# Fast and efficient digestion of adeno associated virus (AAV) capsid proteins for liquid chromatography mass spectrometry (LC-MS) based peptide mapping and post translational modification analysis (PTMs)

**DOI:** 10.1101/2021.08.11.455907

**Authors:** Felipe Guapo, Lisa Strasser, Silvia Millán-Martín, Ian Anderson, Jonathan Bones

**Affiliations:** Characterization and Comparability Laboratory, NIBRT – National Institute for Bioprocessing Research and Training, Foster Avenue, Belfield, Blackrock, Dublin, A94 X099, Ireland; Pharmaron, 12 Estuary Banks, Speke, Liverpool L24 8RB, United Kingdom; School of Chemical and Bioprocess Engineering, University College Dublin, Belfield, Dublin, D04 V1W8, Ireland

**Keywords:** Immobilised pepsin, SMART pepsin digestion, peptide mapping, AAV gene therapy, liquid chromatography mass spectrometry

## Abstract

Adeno-associated virus (AAV) represent a widely used delivery mechanism for gene therapy treatments currently being developed. The size and complexity of these molecules requires the development of sensitive analytical methods for detailed product characterization. Among the quality attributes that need to be monitored, characterization of the AAV capsid protein amino acid sequences and any associated post translational modifications (PTM) present should be performed. As commonly used for recombinant protein analysis, LC-MS based peptide mapping can provide sequence coverage and PTM information to improve product understanding and the development and deployment of the associated manufacturing processes. In the current study, we report a fast and efficient method to digest AAV5 capsid proteins in only 30 minutes prior to peptide mapping analysis. The performance of different proteases in digesting AAV5 was compared and the benefits of using nanoflow liquid chromatography for separation prior to high resolution mass spectrometry to obtain 100% sequence coverage are highlighted. Characterization and quantitation of PTMs on AAV5 capsid proteins when using pepsin as a single protease is reported, thereby demonstrating the potential of this method to aid with complete characterization of AAV serotypes in gene therapy development laboratories.

## 1. Introduction

Adeno-associated virus (AAV) gene therapy products, represent an area of growing interest due to their ability to deliver target genes to human cells, generating long term gene expression with minimal evidence of toxicity or invocation of an immune response [1,2]. However, the biological complexity found in several known AAV serotypes poses a challenge in understanding and optimizing manufacturing processes, increasing the pressure for the development of analytical methods that can perform accurate measurement of capsid protein sequence variability and stability [3].

The icosahedral capsid of AAV is formed by 60 copies of 3 capsid proteins, namely VP1, VP2 and VP3, present in a ratio of approximately 1:1:10, respectively [3]. Although serotype-specific differences in sequence composition have been elucidated previously [4], more information, especially concerning post translational modifications (PTMs), is required. This knowledge could lead to a better understanding of biological differences between serotypes in terms of infectivity, cell targeting and tissue specificity [3,5,6] but also, their molecular stability, which diverges among different serotypes [7–9].

Being able to efficiently characterize VPs is crucial for knowledge-driven improvements of AAV products, such as mixed serotype capsids that can outperform treatments using single serotype AAVs [3,10].

LC-MS based peptide mapping is commonly used in the biopharmaceutical industry to verify the primary structure of a therapeutic protein and to monitor multiple quality attributes which can impact safety and efficacy of the final product. With recent advances in mass spectrometric instrumentation and software platforms, increased accuracy, reproducibility and robustness are now part of efficient peptide mapping workflows [11]. While trypsin is commonly used for proteolytic digestion, it has been found to inefficiently digest VPs of AAV2, requiring alternative proteases, such as Lys C and Asp N, to ensure high sequence coverage [4]. Hence, extensive sample preparation is required, diminishing interest from fast paced pharmaceutical and biotechnological environments.

In this study, we present a semi-automated sample preparation workflow, reducing required hands-on time, while increasing robustness and reliability of generated data. We examined the performance of three proteases, namely, trypsin, chymotrypsin and pepsin, ensuring complete digestion and full sequence coverage of AAV5 capsid proteins in as little as 30 minutes. The chromatographic performance of both, analytical and nano scale LC, and their impact on capsid protein sequence coverage was compared. The resulting workflow demonstrates an easy and fast way to conduct peptide mapping of VP proteins and enabling analysis of PTMs which can ultimately support manufacturing processes and clinical trials of novel AAV-based gene therapy products.

## 2. Materials and Methods

### 2.1 Chemical and reagents

AAV5 empty reference material (06-810) from Applied Viromics (Fremont, CA, USA) was kindly provided by Pharmaron. Thermo Scientific™ SMART Digest™ kits, were obtained from Thermo Fisher Scientific (Sunnyvale, CA, USA). LC-MS grade solvents (0.1% (*v/v*) formic acid (FA) in water, 0.1% (*v/v*) FA in acetonitrile, FA, acetonitrile, water) and Tris (2-carboxyethyl) phosphine hydrochloride (TCEP) were sourced from Fisher Scientific (Dublin, Ireland). All other reagents were purchased from Sigma-Aldrich (Wicklow, Ireland).

### 2.2 Analytical instrumentation

LC-MS analysis was performed using either an UltiMate 3000 RSLCnano system or a Vanquish Flex UHPLC system coupled to a Q Exactive Plus Hybrid quadrupole-Orbitrap mass spectrometer (Thermo Scientific, Bremen, Germany) by means of an EASY-Spray source or by using an Ion Max API source equipped with a heated electrospray ionization (HESI-II) probe (Thermo Fisher Scientific, Bremen, Germany). All data were acquired using Thermo Scientific’s Chromeleon Chromatography Data System software 7.2.9 (Thermo Scientific, Germering, Germany).

### 2.3 Sample preparation for peptide mapping

Four micrograms of AAV5 (4 × 10^11^ vg/mL viral particles) were digested in triplicates using SMART Digest magnetic bead bulk trypsin, chymotrypsin, and pepsin kits. For each digestion, the sample, respective digestion buffer and 5mM TCEP were added to a 96-deepwell plate (Thermo Fisher Scientific, Vantaa, Finland). Samples were incubated for 30 min at 70°C on medium mixing speed to prevent sedimentation of the beads using a Thermo Scientific KingFisher Duo Prime purification system with Thermo Scientific BindIt software version 4.0. At the end of the incubation period, beads were removed, and the remaining volume of the sample was transferred to 1.5 mL Eppendorf protein LoBind tubes (Eppendorf, Dublin) followed by acidification with trifluoroacetic acid (TFA) at a final concentration of 0.1%. Samples were evaporated to dryness and subsequently dissolved in 0.1% FA in water prior to LC-MS analysis.

### 2.4 LC-MS analysis

Resulting peptides were separated using a Thermo Scientific Easy-Spray PepMap RSLC C18 2μm column, 75μm x 50 cm and a Thermo Scientific Hypersil Gold RP, 3 μm, 1mm x 150 mm column (Thermo Fisher, Sunnyvale, CA, USA). Analysis was performed using a binary gradient of 0.1% (*v/v*) FA in water (A) and 0.1% (*v/v)* FA in acetonitrile (B). For analysis, 200 ng and 1μg of digested peptides were injected for each column, respectively.

#### 2.4.1 PepMap RSLC C18 nano LC-MS experimental conditions

Gradient conditions for the Pepmap RSLC C18 were as follows: 2% B to 40% B in 105 min, increased to 60% B in 5 min, increased to 80% in 1 min followed by an isocratic hold for 9 min, followed by 2 linear gradients of 2% to 80% in 5 min, with an isocratic hold for 5 min. Afterwards, the column was re-equilibrated to starting conditions for 20 min. The used flowrate was 250 nL/min and the column temperature was maintained at 45°C.

Data-dependent acquisition (DDA)-MS analysis was performed in positive ion mode. Full scans were acquired at a resolution of 70,000 (at *m/z* 200) for a mass range of *m/z* 200-2,000 using an AGC target of 1.0 × 10^6^ with a maximum injection time (IT) of 100 ms and 1 microscan. Fragment scans were acquired using a resolution setting of 17,500 (at *m/z* 200) with an AGC target of 1.0 × 10^5^ and a maximum IT of 200 ms, an isolation window of *m/z* 2.0 and a signal intensity threshold of 1.0 × 10^4^. Fragmentation of top 10 most abundant precursor ions was done using a normalised collision energy set to 28 using a dynamic exclusion for 45 s and charge exclusion set to unassigned and > 8. MS instrumental tune parameters were set as follows: spray voltage was set to 1.7 kV, capillary temperature was 310°C and S-lens RF voltage set to 50.

#### 2.4.2 Hypersil Gold RP LC-MS experimental conditions

Gradient conditions for the Hypersil Gold RP were as follows: 2% B to 40% B in 105 min, increase to 60% B in 5 min, increase to 80% in 1 min followed by an isocratic hold for 14 min. Column was re-equilibrated at 2% B for 10 min at flow rate of 100 μL/min and column temperature was maintained at 60°C.

Data-dependent acquisition (DDA)-MS analysis of top 10 most abundant ions was performed in positive ion mode as above apart from the AGC target which was set to 3.0 × 10^6^ and the dynamic exclusion time which was set to 7 s. MS instrumental tune parameters were set as follows: spray voltage was set to 3.8 kV, sheath gas flow rate was 25 arbitrary units (AU), auxiliary gas flow rate was 10 AU, capillary temperature was 320°C, probe heater temperature was 150°C and S-lens RF voltage set to 60.

### 2.5 Data processing

Peptide identification and PTM relative quantitation were performed using the Thermo Scientific BioPharma Finder software version 4.1 (Thermo Scientific, San Jose, CA, USA) using parameters summarised in Table S1. The results were filtered in two stages: 1) for sequence coverage evaluation and 2) for PTMs relative quantitation. The list of identified peptides was filtered to exclude nonspecific generated peptides, unknown modifications and gas phase ions. Peptide lists included all the peptides within ± 5 ppm accuracy, 1 × 10^5^ average MS area and a confidence score ≥ 95%.

The sequence coverage percentage for each VP protein is reported for each experimental condition and a final list of evaluated PTMs with relative abundances ≥ 1% are reported for the pepsin digested samples.

## 3. Results

### 3.1 Fast and efficient digestion of AAV capsid proteins

In the present study, a sample preparation workflow for peptide mapping of AAV capsid proteins was optimized comparing different commonly used proteases, trypsin, chymotrypsin, and pepsin. Employing enzymes immobilised on magnetic beads allowed for automation, resulting in minimized hands-on time while yielding in excellent reproducibility in only 30 minutes.

An assessment of the SMART Digest trypsin kit following a peptide mapping protocol commonly used for monoclonal antibodies (mAbs) as reported by Millán Martín *et al*. [12] was conducted to gather preliminary results regarding digestion efficiency for AAV5. The results of this analysis showed insufficient digestion of AAVs with a sequence coverage below 50% (data not shown), clearly highlighting the need to improve sample preparation conditions. Furthermore, the generally low concentration of AAV samples led to the conclusion that higher sensitivity for LC-MS analysis would be beneficial. Hence, the use of nanoflow LC separation was explored.

Pepsin has low specificity with multiple cleavage sites; thus, offering a better chance to fully digest AAV capsid proteins. However, reproducibility is often noted as being problematic [13,14]. The Thermo SMART Digest kit aims at resolving this problem, by removing the protease from the sample thereby rapidly stopping the reaction in a controlled manner resulting in highly reproducible results.

Figure 1 shows the base peak chromatogram (BPC) obtained from peptide mapping experiments using the PepMap RSLC C18 column, which highlights how pepsin was able to more efficiently digest AAV5, producing populated chromatograms showing a larger population of peptide peaks while also covering areas not addressed by either trypsin or chymotrypsin.

**Figure 1:**
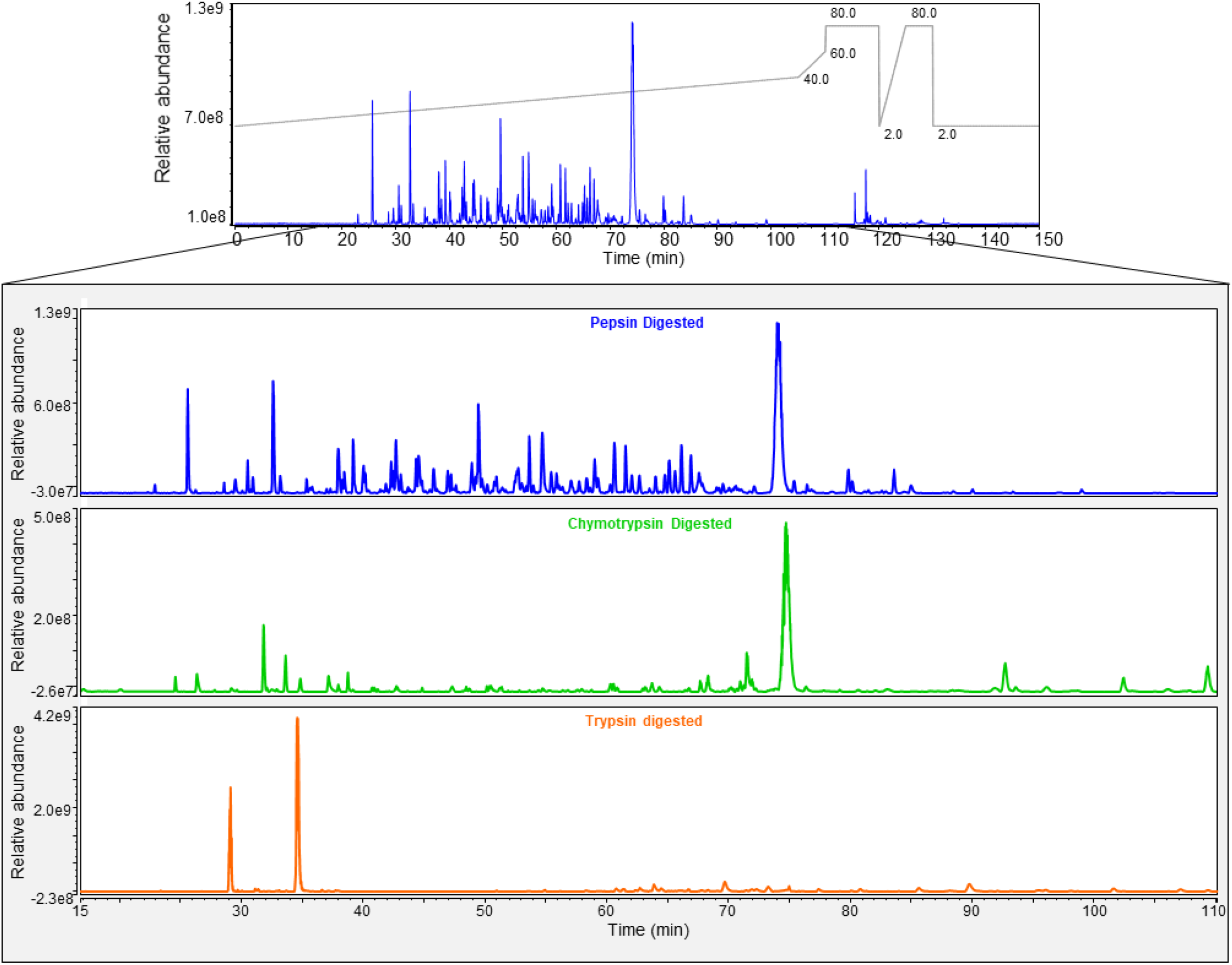
(Top) Base peak chromatogram (BPC) obtained from peptide mapping experiment using pepsin digestion resolved on a PepMap RSLC C18 column. Indicated in grey is the acetonitrile-gradient used for separation. (Bottom) Zoomed view (15 – 110 min) of BPCs obtained from peptide mapping experiments of AAV5 for the digestion experiment using pepsin (blue), chymotrypsin (green) and trypsin (orange).

To ensure reliability of results, digestions were performed in triplicate. The total sequence coverage was evaluated for all three capsid proteins of AAV5 comparing the different proteases (Table 1). Our data demonstrates that pepsin provides a fast and efficient digestion of AAV5 capsid proteins, resulting in a sequence coverage of 100% for all three VPs in just 30 min of digestion (Table 1). This represents a considerable reduction of the required overnight incubation time reported in Rumachik et al. [15] without the need of additional sample preparation steps.

**Table 1:** Sequence coverage and total VP1 assigned MS peaks for each replicate and protease after 30 min digestion resolved on a PepMap RSLC C18 column.

| Protease | MS Peaks |  |  |  |
| --- | --- | --- | --- | --- |
|  | VP1 | Sequence coverage (%) |  |  |
|  |  | VP1 | VP2 | VP3 |
| Pepsin 1 | 1637 | 100 | 100 | 100 |
| Pepsin 2 | 1653 | 100 | 100 | 100 |
| Pepsin 3 | 1593 | 100 | 100 | 100 |
| Trypsin 1 | 18 | 30.9 | 24.5 | 25.6 |
| Trypsin 2 | 18 | 30.9 | 24.5 | 25.6 |
| Trypsin 3 | 18 | 30.9 | 24.5 | 25.6 |
| Chymotrypsin 1 | 09 | 16 | 19.7 | 30.1 |
| Chymotrypsin 2 | 09 | 16 | 19.7 | 30.1 |
| Chymotrypsin 3 | 09 | 16 | 19.7 | 30.1 |

In comparison to pepsin, trypsin and chymotrypsin were not able to sufficiently digest the capsid proteins with sequence coverages of VP1 falling below 30.9 and 16%, respectively (Table 1 and Figure 2). Pepsin requires a much lower pH than the other two proteases to be active. Thus, the low pH used in the digestion buffer is potentially aiding capsid protein denaturation and unfolding which exposes the cleavage sites, ensuring complete digestion.

**Figure 2:**
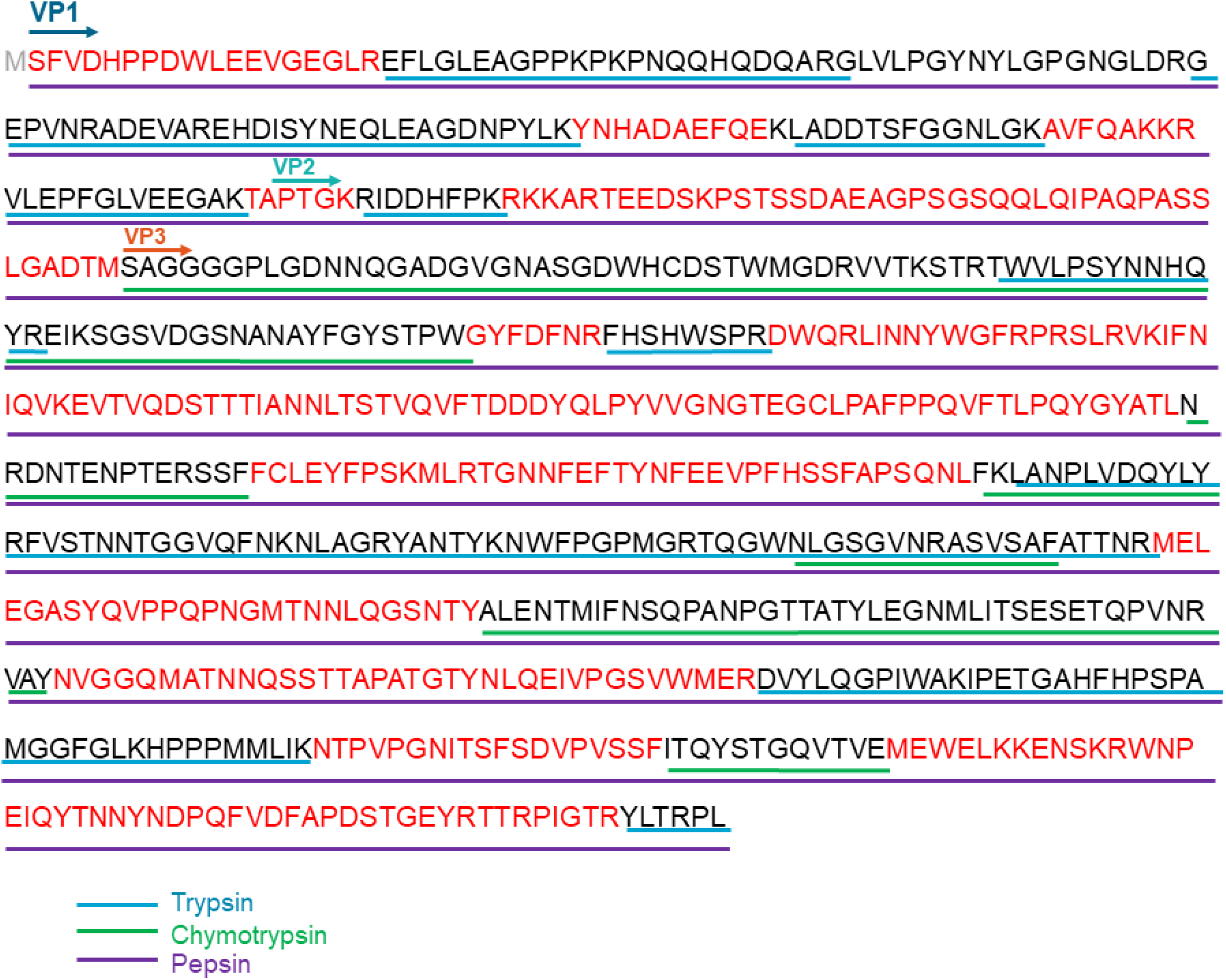
Sequence coverage map of AAV5 capsid proteins (VP1, 2 and 3) by LC-MS/MS using different enzymatic digests. Green line: chymotryptic peptides, blue line: tryptic peptides, purple line: pepsin peptides. Protein sequence only identified by pepsin highlighted in red.

To ease visualization in BPF a filter of ≥ 1% minimum peptide recovery was applied reducing the number of peptides displayed in the sequence coverage map for pepsin (Figure S1). In BPF the recovery value displays the relative abundance of the modified peptide normalized against the peak areas of all peptides. Meaning higher-intensity peptides score more than lower-intensity peptides. Typically, a fair recovery is greater than 1%.

It is important to note that with this filter, sequence coverage was reduced to 96.5, 100 and 100% for VP1, VP2 and VP3, respectively, as two peptides in the N terminal of VP1 are only confirmed by peptides of less than 1% recovery (Figure S1).

### 3.2 Quantitative analysis of PTMs in AAV5

Relative levels of PTMs were studied for the pepsin digested and PepMap RSLC C18 resolved VP peptides only, as these were the only ones to achieve 100% sequence coverage. PTM signatures were defined based on the differential (delta) mass values of modified *versus* unmodified peptides generated by pepsin digestion in any of the screened samples and identified automatically by BPF. Manual validation was performed in addition to software assignment to increase PTM identification confidence. Variable modifications were monitored as shown in Table S1. Importantly, using this method did not provide a specific capsid protein location for PTMs, as most of the VP1 sequence is shared with VP2 and VP3.

Relative levels of each PTM were studied and quantified using triplicate sample analysis. After filtering the results as described in section 2.5, a total of 21 PTMs with at least 1% relative abundance were reported, with 5 sites identified as containing deamidations and 6 containing oxidations. In addition to deamidations and oxidations, 2 acetylation, 1 methylation, 2 isomerization, 1 amidation and 4 succinimide formation are reported with relative abundance values above 1% (Table 2).

**Table 2:**
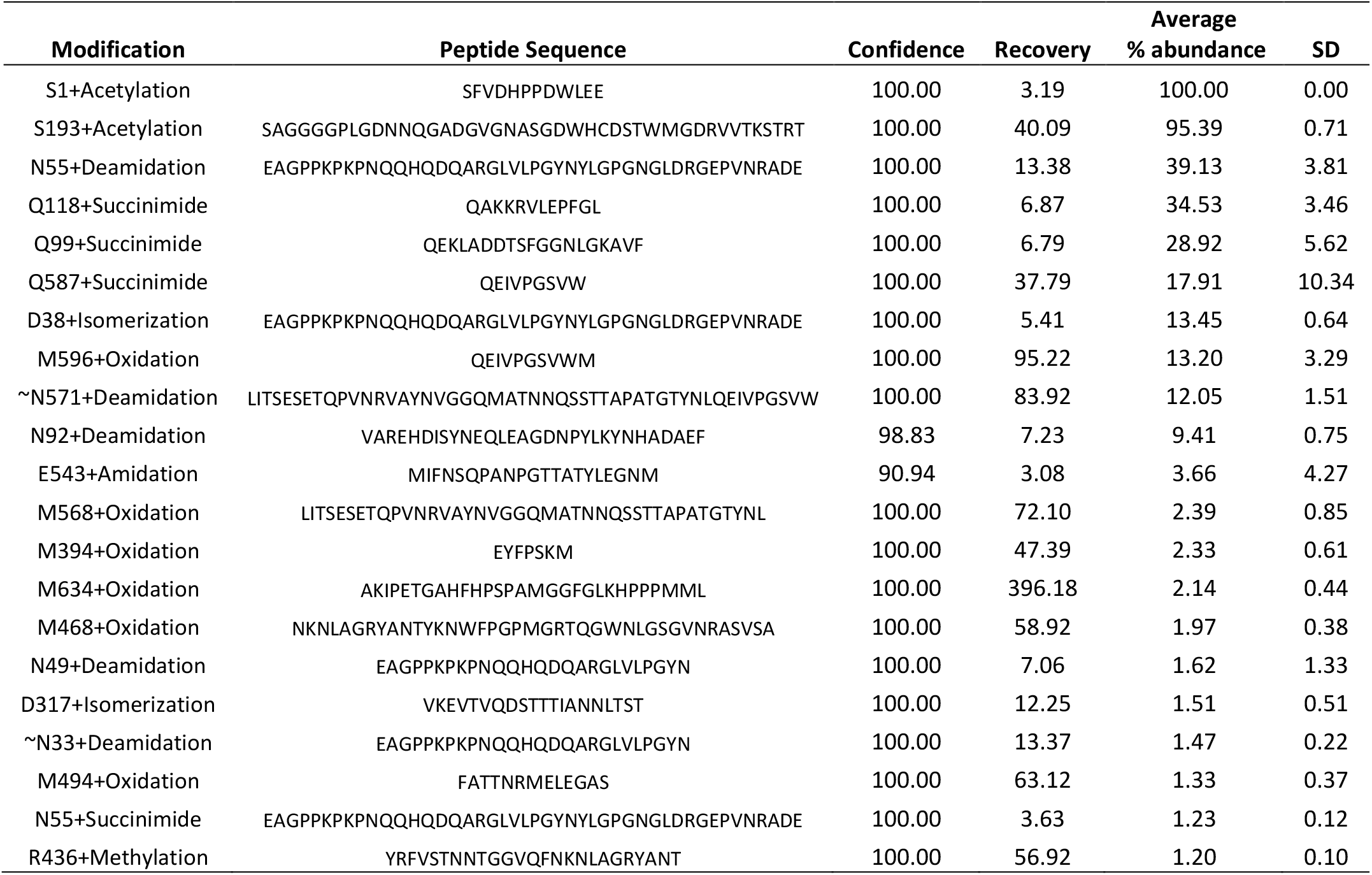
Summary of PTMs identified, resolved on a PepMap RSLC C18 column and quantified for AAV5 with ≥ 1% relative abundance. Modifications marked with a tilde (~) indicate the approximate location of the modification as BPF was unable to determine the exact modification site in that instance.

| Modification | Peptide Sequence | Confidence | Recovery | Average % abundance | SD |
| --- | --- | --- | --- | --- | --- |
| S1+Acetylation | SFVDHPPDWLEE | 100.00 | 3.19 | 100.00 | 0.00 |
| S193+Acetylation | SAGGGGPLGDNNQGADGVGNASGDWHCDSTWMGDRVVTKSTRT | 100.00 | 40.09 | 95.39 | 0.71 |
| N55+Deamidation | EAGPPKPKPNQQHQDQARGLVLPGYNYLGPNGLDRGEPVNRAD | 100.00 | 13.38 | 39.13 | 3.81 |
| Q118+Succinimide | QAKKRVLEPFGL | 100.00 | 6.87 | 34.53 | 3.46 |
| Q99+Succinimide | QEKLADDTSFGGNLGKAVF | 100.00 | 6.79 | 28.92 | 5.62 |
| Q587+Succinimide | QEIVPGSVW | 100.00 | 37.79 | 17.91 | 10.34 |
| D38+Isomerization | EAGPPKPKPNQQHQDQARGLVLPGYNYLGPNGLDRGEPVNRAD | 100.00 | 5.41 | 13.45 | 0.64 |
| M596+Oxidation | QEIVPGSVWM | 100.00 | 95.22 | 13.20 | 3.29 |
| ~N571+Deamidation | LITSESETQPVNRVAYNVGGQMATNNQSSTTAPATGTYNLQEIVPGSVW | 100.00 | 83.92 | 12.05 | 1.51 |
| N92+Deamidation | VAREHDISYNEQLEAGDNPYLKYNHADA | 98.83 | 7.23 | 9.41 | 0.75 |
| E543+Amidation | MIFNSQPANPGTTATYLEGNM | 90.94 | 3.08 | 3.66 | 4.27 |
| M568+Oxidation | LITSESETQPVNRVAYNVGGQMATNNQSSTTAPATGTYNL | 100.00 | 72.10 | 2.39 | 0.85 |
| M394+Oxidation | EYFPSKM | 100.00 | 47.39 | 2.33 | 0.61 |
| M634+Oxidation | AKIPETGAHFHPSAMGGFGLKHPPMML | 100.00 | 396.18 | 2.14 | 0.44 |
| M468+Oxidation | NKNLAGRYANTYKNWFGPMGRTQGWNLGSGVNRASVSA | 100.00 | 58.92 | 1.97 | 0.38 |
| N49+Deamidation | EAGPPKPKPNQQHQDQARGLVLPGYN | 100.00 | 7.06 | 1.62 | 1.33 |
| D317+Isomerization | VKEVTVDSTTTIANNLST | 100.00 | 12.25 | 1.51 | 0.51 |
| ~N33+Deamidation | EAGPPKPKPNQQHQDQARGLVLPGYN | 100.00 | 13.37 | 1.47 | 0.22 |
| M494+Oxidation | FATTNRMELEGAS | 100.00 | 63.12 | 1.33 | 0.37 |
| N55+Succinimide | EAGPPKPKPNQQHQDQARGLVLPGYNYLGPNGLDRGEPVNRAD | 100.00 | 3.63 | 1.23 | 0.12 |
| R436+Methylation | YRFVSTNNTGGVQFNKNLAGRYANT | 100.00 | 56.92 | 1.20 | 0.10 |

The top 10 most abundant modifications including S1 and S193 acetylation; N55, N92 and ~N571 deamidation; Q99, Q118 and Q587 succinimide formation; D38 isomerization and M596 oxidation are displayed in figure 3. The MS/MS spectrum and peptide fragment coverage map for the top 5 PTMs are included in the supplementary material. A complete list of all 72 identified PTMs including low abundant modifications (< 1% relative abundance) is also listed in table S2.

**Figure 3:**
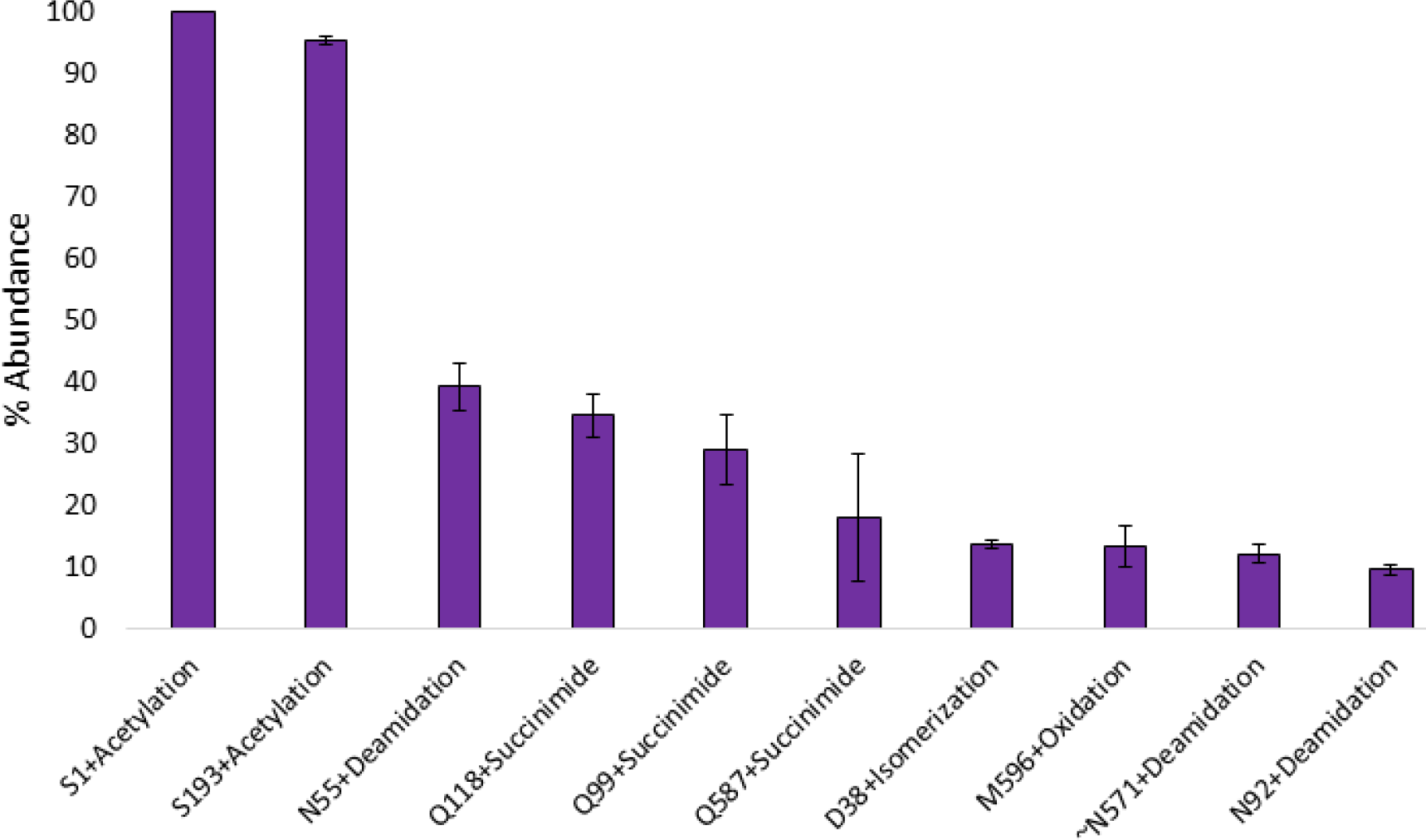
Average relative abundance (n = 3) of top 10 identified PTMs in AAV5 empty reference material digested with pepsin (30 min) and resolved on a PepMap RSLC C18 column.

### 3.3 LC-MS/MS analysis on the Hypersil Gold RP column using an analytical flow rate

To test the impact on chromatographic sensitivity and the possibility of using a more robust workflow for the identification of AAV peptides and PTMs, the same samples digested with pepsin were analysed using analytical flow LC separation (100 μL/min).

Thereby, sequence coverage was found to be reduced to 91.7, 91.3 and 100% for VP1, VP2 and VP3 respectively (Table 3) indicating a loss in sensitivity when using the higher flow rate, which also impacted PTM identification. Only 32 PTMs were identified using the Hypersil Gold RP even though 5 times more sample was injected for this analysis (Table S3).

**Table 3:**
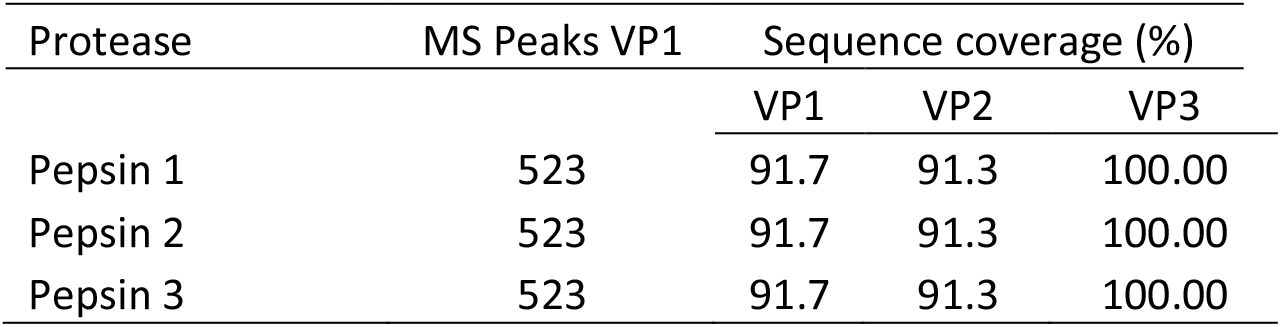
Sequence coverage for each replicate digested with pepsin and run on a Hypersil Gold RP at 100 μL/min.

| Protease | MS Peaks VP1 | Sequence coverage (%) |  |  |
| --- | --- | --- | --- | --- |
|  |  | VP1 | VP2 | VP3 |
| Pepsin 1 | 523 | 91.7 | 91.3 | 100.00 |
| Pepsin 2 | 523 | 91.7 | 91.3 | 100.00 |
| Pepsin 3 | 523 | 91.7 | 91.3 | 100.00 |

## 4. Discussion

A quick and efficient peptide mapping protocol for AAV-based therapeutic products is key for informing upstream development processes regarding identifying amino acid sequence changes and PTMs, with the potential to affect product stability and efficiency.

The presented workflow demonstrates a significant improvement compared to previously reported sample preparation methods which may take up to 18 hours while also requiring larger amounts AAVs to produce satisfactory results [15]. Additionally, the number of manipulation steps required in traditional and more recent digestion protocols was also reduced [16]. By using a magnetic bead-based sample preparation for the digestion with pepsin, digestion is stopped in all samples at the exact same time, avoiding the production of different digestion patterns. Thereby, reproducibility is increased while also taking advantage of pepsin’s lower specificity in comparison to trypsin, to produce quick results. Furthermore, pepsin has the advantage of being active at low pH. This aids the breakdown of the viral particles in association with the high temperatures used in the presented digestion protocol, helping to overcome the high thermal stability possessed by AAV5 that may deter protein proteolysis [17], ensuring digestion efficiency and high sequence coverage in a short amount of time.

Using the presented workflow resulted in 100% sequence coverage as well as the identification of 72 PTMs, 21 of each are confidently quantified with relative abundances above 1%. These represent quality attributes that can inform process development. While digestion with pepsin was carried out at low pH, which is known to reduce deamidation incidence [18], 2 sites (N55 and ~N571) had relative abundances above 10%. The high temperature (70°C) used for the digestion could contribute to increasing deamidation levels. However, a similar workflow was not seen to increase deamidation levels in mAbs [12]. Therefore, further investigations are necessary, to study the actual abundance of process induced deamidation in AAV samples.

Oxidation is also a PTM of interest to assess process induced variations. All 16 oxidation events detected had a relative abundance of below 2.5% except for M596. Oxidation can be a result of conditions encountered during purification, formulation and analysis as well as prolonged storage and frequent freeze-thaw cycles. Hence, the presence of oxidation constitutes a critical quality attribute, as oxidation can adversely impact the activity and stability of biotherapeutics [19].

Other PTMs like isomerization and succinimide formation are also of interest as they can alter protein structure and function, influencing binding and overall performance of the pharmacological product [20,21]. These processes can occur *in vivo* and *in vitro* and are influenced by environmental conditions. Thus, their monitoring is of pharmacological importance.

When it comes to AAV-based gene therapy products, limited information is available regarding PTM levels across different serotypes and the biological implication of such PTMs. Employing intact protein analysis, Jin *et al.* [4] demonstrated that the first methionine of VP1 and 3 is cleaved off after synthesis, and the second amino acid of the sequence is subsequently acetylated. Using the presented peptide mapping workflow allowed for successful detection and relative quantitation of an acetylation on the suggested amino acid. Furthermore, Mary *et al.* [6] describes the detection of 9 different PTMs for AAV5 after tryptic digestion of AAV-293 cell generated viral capsids, including 3 ubiquination events, 1 phosphorylation, 1 SUMOylation and 4 glycosylation events. Our dataset confirms the identification of phosphorylation at S679 (S680 when the M at position 1 is considered), but the remaining modifications identified could not be confirmed using the present dataset. More recently, acetylation, deamidation, oxidation, methylation, and phosphorylation modifications were also identified in AAV5 after a 98% successful tryptic digestion [16]. General discrepancies on PTM abundance levels can be explained by differences in expression systems or overall downstream processing. Nevertheless, it is important to highlight that our workflow using pepsin digestion provides 100% sequence coverage in 30 min, giving a better insight into AAV5 sequence and associated PTMs.

In addition, the challenge posed by low concentrated AAV samples means that limited amounts of the biotherapeutic product are usually available for analysis. Thus, sensitivity becomes a critical parameter as demonstrated by our comparison between the use of analytical or nanoflow LC prior to MS detection. A trade off needs to be considered regarding sample availability and robustness, when deciding on what system to run the analysis on.

Moreover, pepsin poses its own challenges that need to be examined during data analysis. Due to its low specificity, applying a missed cleavage filter, which is commonly used for tryptic digests was deemed not appropriate. As smaller peptides are generated, an impact on peptide intensity spectra was also observed, which makes the data analysis challenging. Thus, it was decided to only highlight PTMs found with at least 1% relative abundance. However, a list of all identified PTMs is reported in table S2.

## 5. Conclusion

The results reported in the present study demonstrate a complete and efficient peptide mapping workflow to successfully monitor the amino acid sequence as well as PTMs of AAV capsid proteins (VPs). The aim of this work was to highlight the advantage of using pepsin over other more traditional proteases such as trypsin and chymotrypsin for peptide mapping analysis of AAVs. Additionally, it was shown that the use of nanoflow chromatography yielded superior sensitivity resulting in an increased sequence coverage as well as improved detection of low abundant PTMs.

Pepsin has been demonstrated to provide access to previously inaccessible areas of the AAV sequence generating fast and precise digestions in 30 minutes, that are easily controlled and stopped by removal of the enzyme at the end of the specified digestion time ensuring high reproducibility. We anticipate that our workflow will greatly benefit fast paced pharmaceutical environments that aim to quickly and completely characterize AAV based gene therapy products through peptide mapping analysis.

## Acknowledgment

The authors greatly acknowledge funding from Enterprise Ireland under the Innovation Partnership Program IP/2018/0753 and support from Abbvie and Pharmaron.

## Author statement

**Felipe Guapo:** Conceptualization, Methodology, Validation, Formal analysis, Investigation, Writing - Original Draft, Visualization. **Lisa Strasser:** Conceptualization, Writing - Review & Editing, Visualization, Supervision. **Silvia Millán-Martín:** Validation, Writing - Review & Editing. **Ian Anderson:** Writing - Review & Editing, Project administration. **Jonathan Bones:** Conceptualization, Supervision, Project administration, Writing - Review & Editing, Funding acquisition.

All authors have given approval to the final version of the manuscript.

## Appendix A. Supplementary material

Supplementary material related to this article can be found below.

## Appendix A. Supplementary Material

**Table S1:**
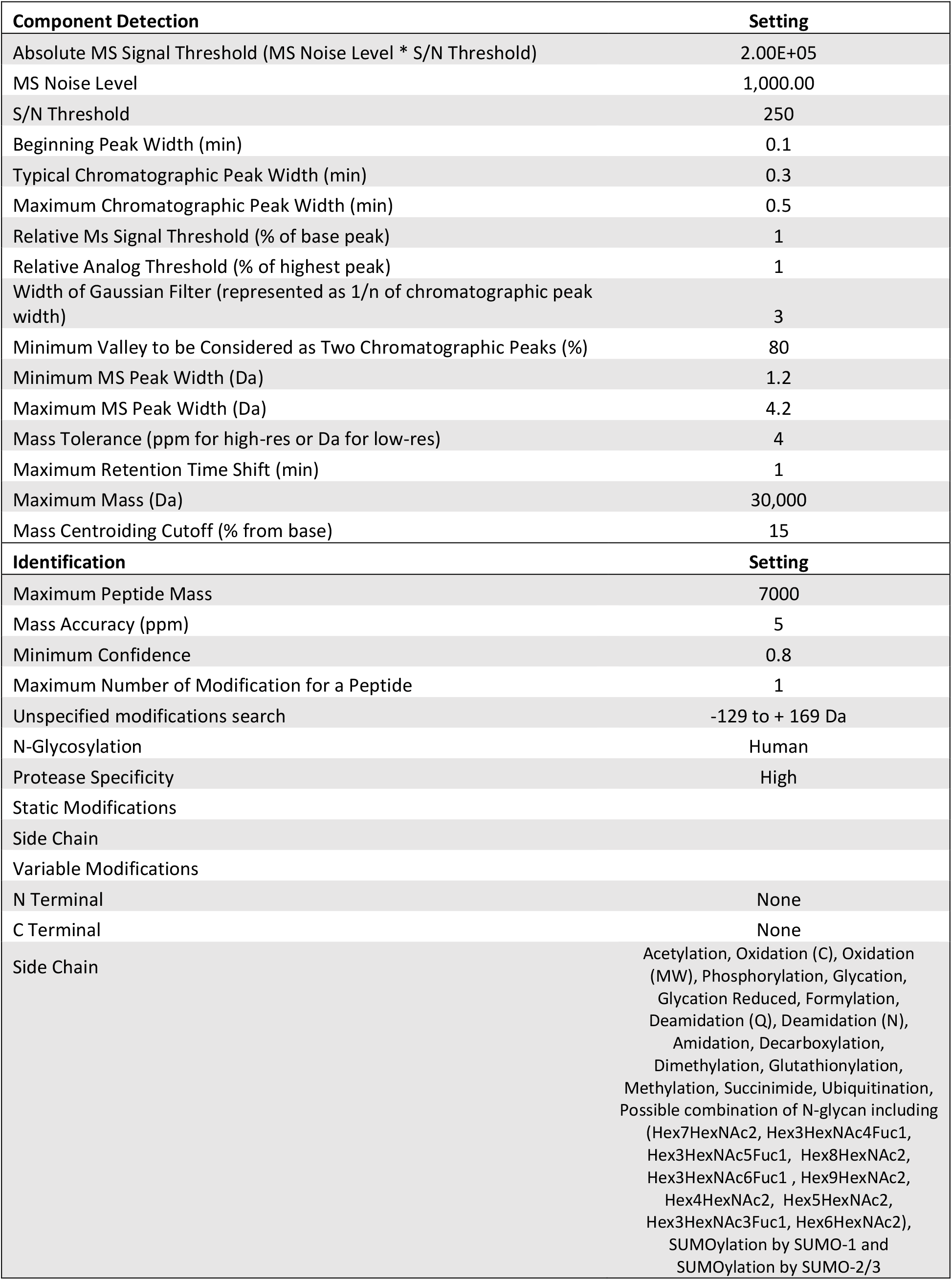
BPF 4.1 software parameter settings for peptide mapping data analysis.

| Component Detection | Setting |
| --- | --- |
| Absolute MS Signal Threshold (MS Noise Level * S/N Threshold) | 2.00E+05 |
| MS Noise Level | 1,000.00 |
| S/N Threshold | 250 |
| Beginning Peak Width (min) | 0.1 |
| Typical Chromatographic Peak Width (min) | 0.3 |
| Maximum Chromatographic Peak Width (min) | 0.5 |
| Relative Ms Signal Threshold (% of base peak) | 1 |
| Relative Analog Threshold (% of highest peak) | 1 |
| Width of Gaussian Filter (represented as 1/n of chromatographic peak width) | 3 |
| Minimum Valley to be Considered as Two Chromatographic Peaks (%) | 80 |
| Minimum MS Peak Width (Da) | 1.2 |
| Maximum MS Peak Width (Da) | 4.2 |
| Mass Tolerance (ppm for high-res or Da for low-res) | 4 |
| Maximum Retention Time Shift (min) | 1 |
| Maximum Mass (Da) | 30,000 |
| Mass Centroiding Cutoff (% from base) | 15 |
| Identification | Setting |
| Maximum Peptide Mass | 7000 |
| Mass Accuracy (ppm) | 5 |
| Minimum Confidence | 0.8 |
| Maximum Number of Modification for a Peptide | 1 |
| Unspecified modifications search | -129 to + 169 Da |
| N-Glycosylation | Human |
| Protease Specificity | High |
| Static Modifications |  |
| Side Chain |  |
| Variable Modifications |  |
| N Terminal | None |
| C Terminal | None |
| Side Chain | Acetylation, Oxidation (C), Oxidation (MW), Phosphorylation, Glycation, Glycation Reduced, Formylation, Deamidation (Q), Deamidation (N), Amidation, Decarboxylation, Dimethylation, Glutathionylation, Methylation, Succinimide, Ubiquitination, Possible combination of N-glycan including (Hex7HexNAc2, Hex3HexNAc4Fuc1, Hex3HexNAc5Fuc1, Hex8HexNAc2, Hex3HexNAc6Fuc1, Hex9HexNAc2, Hex4HexNAc2, Hex5HexNAc2, Hex3HexNAc3Fuc1, Hex6HexNAc2), SUMOylation by SUMO-1 and SUMOylation by SUMO-2/3 |

**Figure S1:**
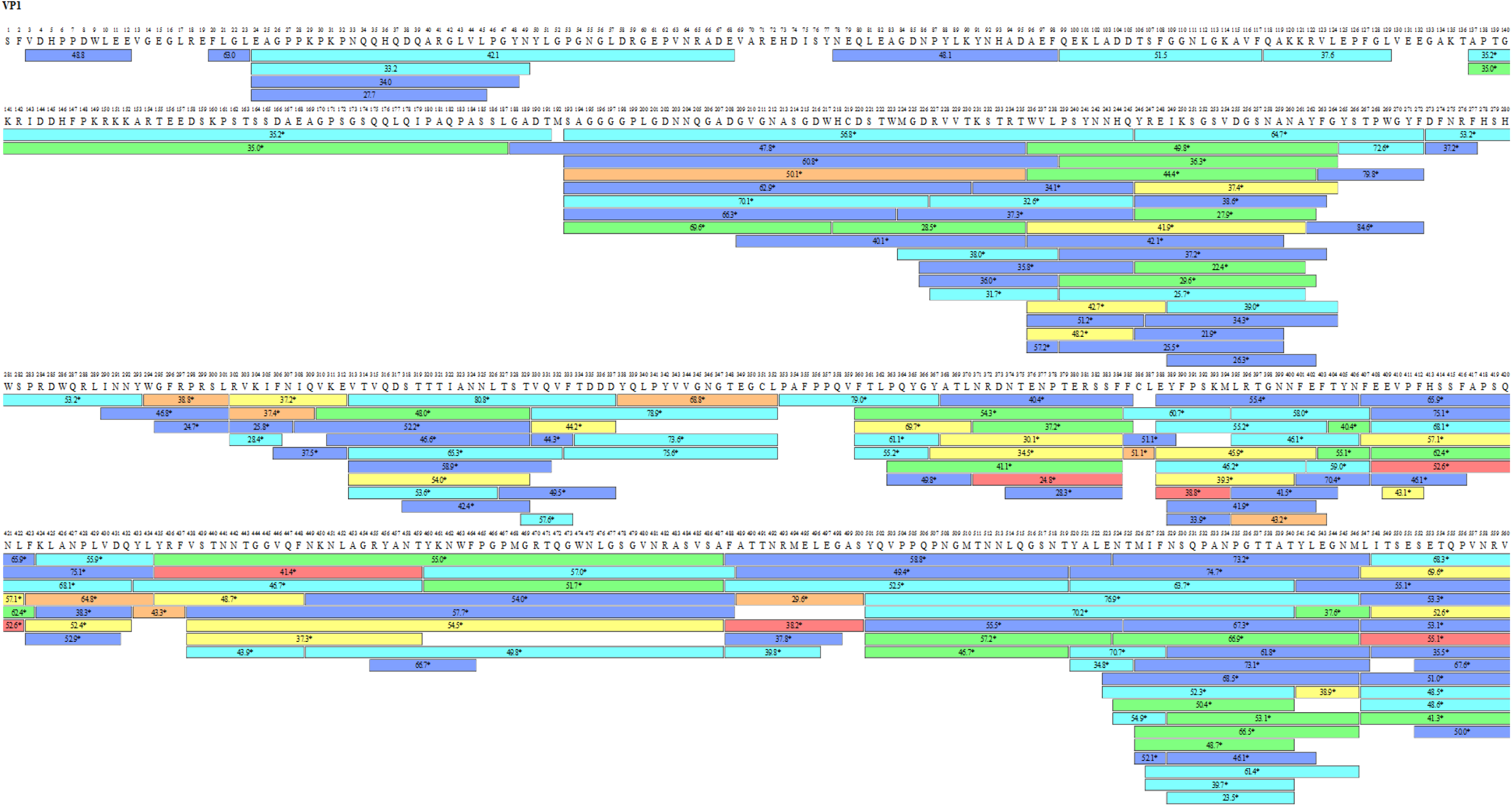
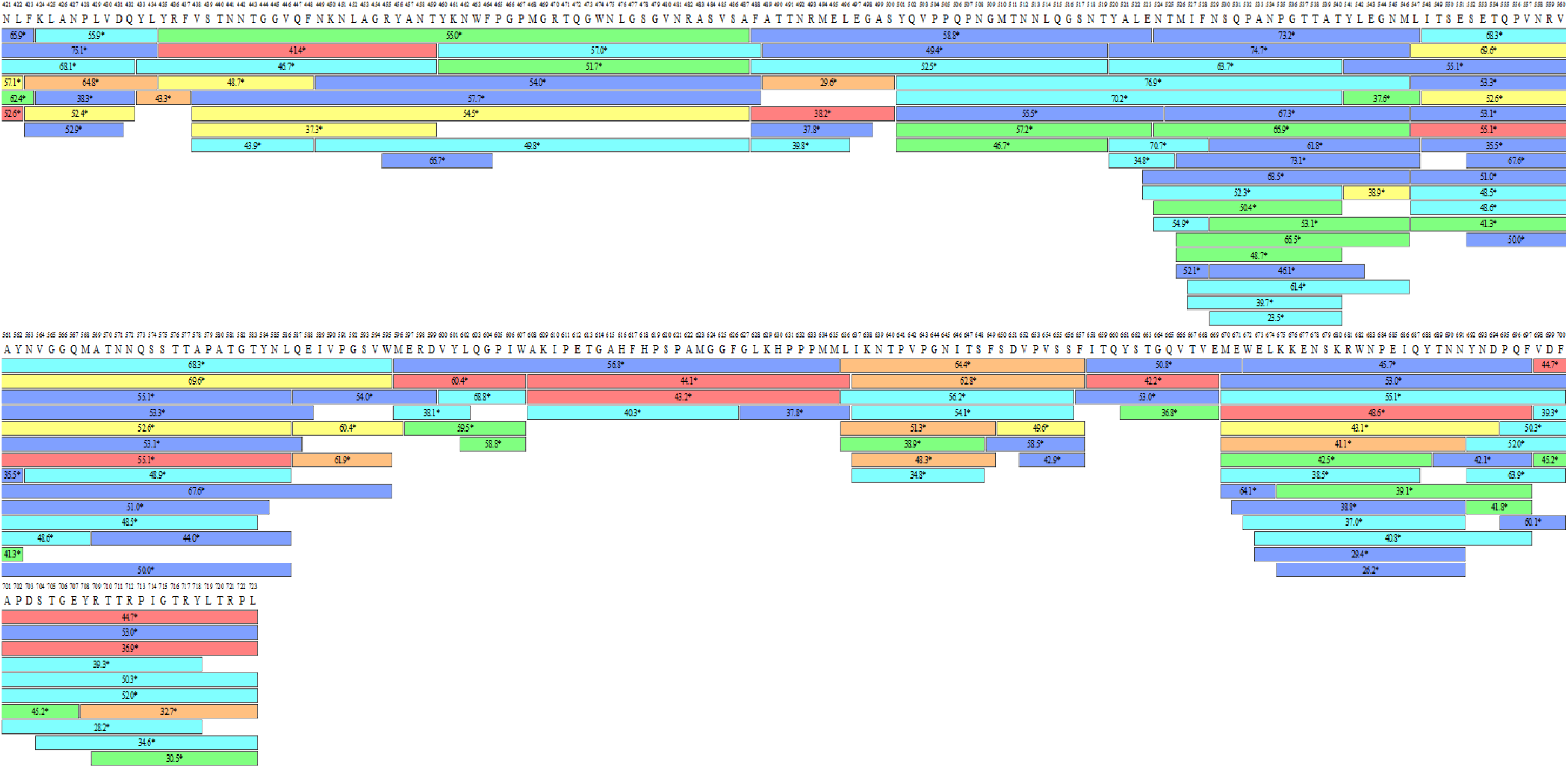
Pepsin sequence coverage map for AAV5 capsid proteins on the PepMap nano RSLC C18 column. The coloured bars show the identified peptides, with the numbers in the bars reflecting the retention time. The different colours indicate the peptide recovery in the MS1 scan: red >50%, orange >20% and yellow >10% represent good recovery. Green, >5%, light blue >2% and cyan >1% represent fair recovery.

**Table S2:**
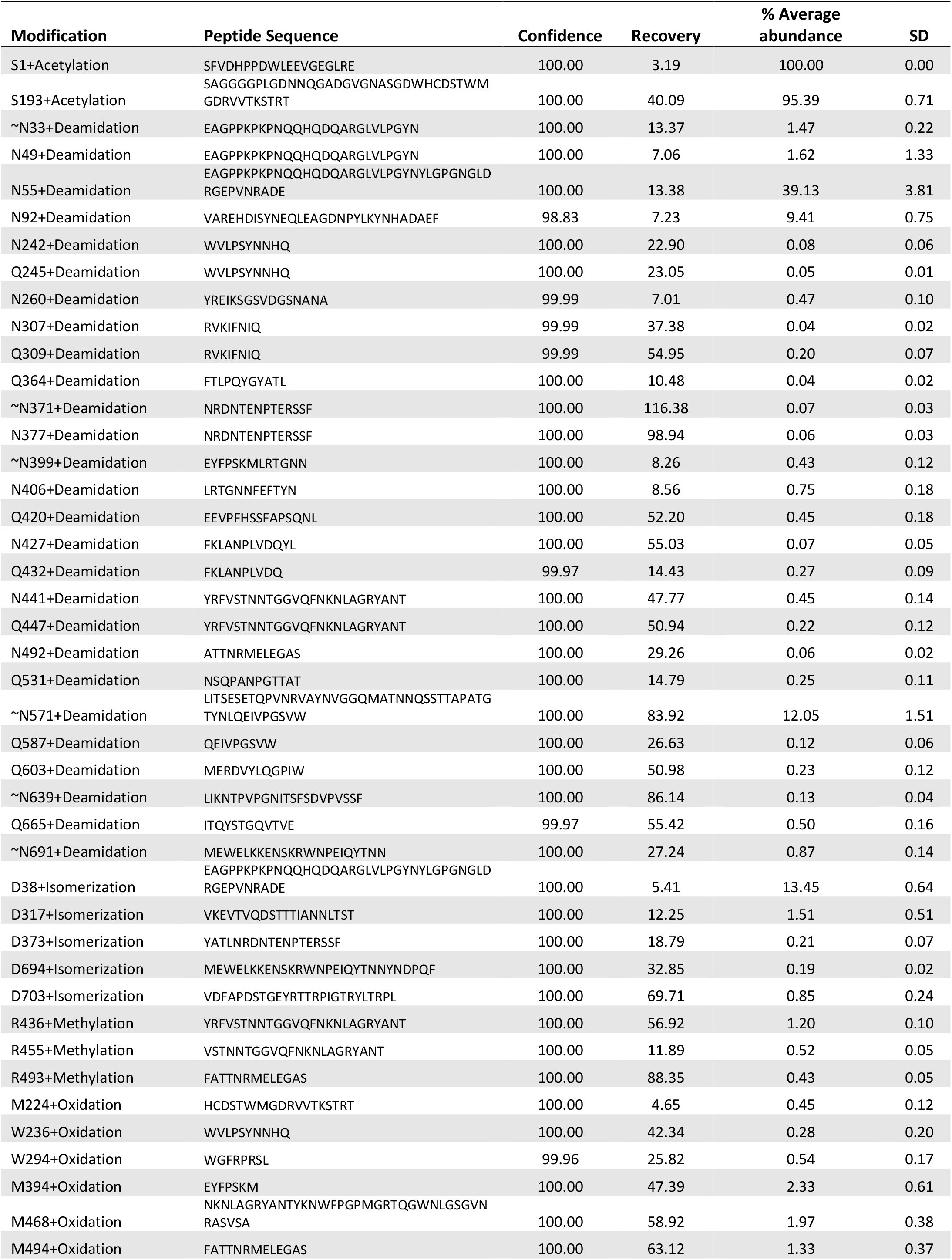
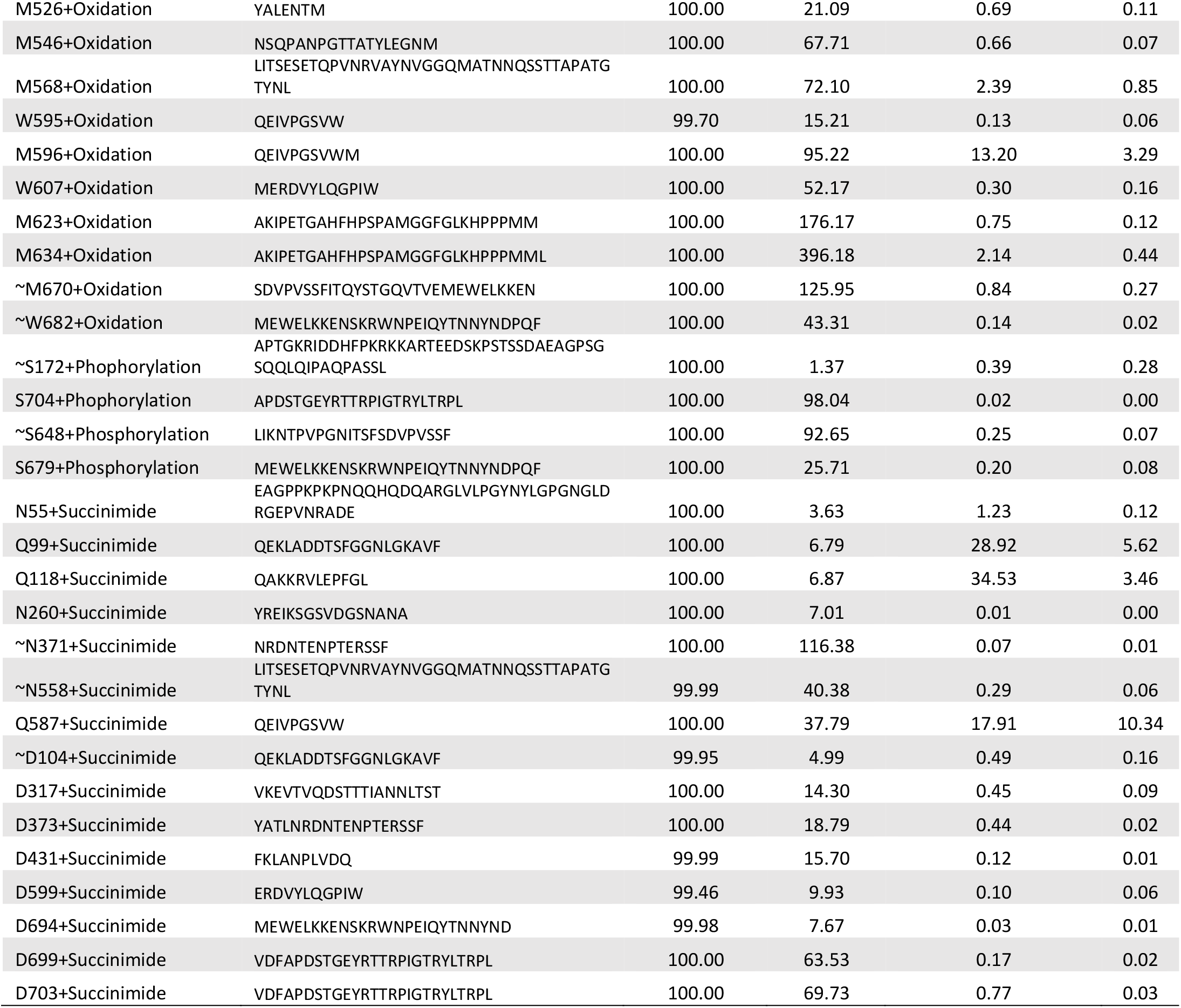
Complete list of PTMs identified, resolved on a PepMap RSLC C18 column and quantified for AAV5.

**Table S3:** Complete list of PTMs identified for AAV5 on Hypersil Gold RP column.

| Modification | Peptide Sequence | Confidence | Recovery |
| --- | --- | --- | --- |
| N55+Deamidation | EAGPPKPKPNQQHQDQARGLVLPGYNYLGPNGLDRGEPVNRAD | 100.00 | 4.45 |
| Q245+Deamidation | WVLPSTNNHQ | 100.00 | 43.59 |
| ~N400+Deamidation | EYFSPKMLRTGNN | 100.00 | 17.04 |
| N406+Deamidation | LRTGNNFEFTYN | 100.00 | 8.50 |
| Q420+Deamidation | EEVPFHSSFAPSQNL | 100.00 | 127.90 |
| Q432+Deamidation | FKLANPLVDQYL | 100.00 | 125.15 |
| ~N571+Deamidation | LITSESETQPVNRVAYNVGGQMATNNQSSTTAPATGTYNL | 100.00 | 142.03 |
| Q603+Deamidation | MERDVYLQGPW | 100.00 | 100.17 |
| ~Q660+Deamidation | ITQYSTGQVTVE | 100.00 | 87.68 |
| ~N691+Deamidation | MEWELKKENSKRWNPEIQYTNN | 100.00 | 41.41 |
| ~Q13+Deamidation | SAGGGGPLGDNNQGADGVGNASGDWHCDSTWMGDRVVTKSTRT | 100.00 | 3.77 |
| D317+Isomerization | VTVDSTTTIANNLTST | 100.00 | 23.05 |
| R436+Methylation | YRFVSTNNTGGVQFNKNLAGRYANT | 100.00 | 66.20 |
| W236+Oxidation | WVLPSTNNHQ | 100.00 | 45.02 |
| M394+Oxidation | EYFSPKM | 100.00 | 75.89 |
| M468+Oxidation | YKNWFPGPMGRTQGWNLGSGVNRASVSA | 100.00 | 136.52 |
| M494+Oxidation | FATTNRMELEGAS | 100.00 | 84.55 |
| M568+Oxidation | LITSESETQPVNRVAYNVGGQMATNNQSSTTAPATGTYNL | 100.00 | 171.49 |
| M596+Oxidation | MERDVYLQGPW | 100.00 | 130.45 |
| W607+Oxidation | MERDVYLQGPW | 99.99 | 102.82 |
| M623+Oxidation | AKIPETGAHFHSPAMGGFGLKHPPPM | 100.00 | 276.82 |
| M634+Oxidation | AKIPETGAHFHSPAMGGFGLKHPPPM | 100.00 | 289.88 |
| M635+Oxidation | AKIPETGAHFHSPAMGGFGLKHPPPMML | 100.00 | 404.38 |
| ~W682+Oxidation | MEWELKKENSKRWNPEIQYTNNYNDPQF | 100.00 | 61.50 |
| ~M32+Oxidation | SAGGGGPLGDNNQGADGVGNASGDWHCDSTWMGDRVVTKSTRT | 100.00 | 5.77 |
| Q99+Succinimide | QEKLADDTSGGNLGKAVF | 100.00 | 12.48 |
| Q118+Succinimide | QAKKRVLEPFGL | 100.00 | 9.15 |
| Q587+Succinimide | QEIVPGSVW | 100.00 | 57.91 |
| D373+Succinimide2 | NRDNTENPTERSF | 100.00 | 117.32 |
| D703+Succinimide2 | VDFAPDSTGEYRTTRPIGTRYLTRPL | 100.00 | 178.60 |
| D16+Succinimide2 | SAGGGGPLGDNNQGADGVGNASGDWHCDSTWMGDRVVTKSTRT | 99.46 | 6.58 |
| N20+Unglycosylated | SAGGGGPLGDNNQGADGVGNASGDWHCDSTWMGDRVVTKSTRT | 100.00 | 126.81 |

**Figure S2:**
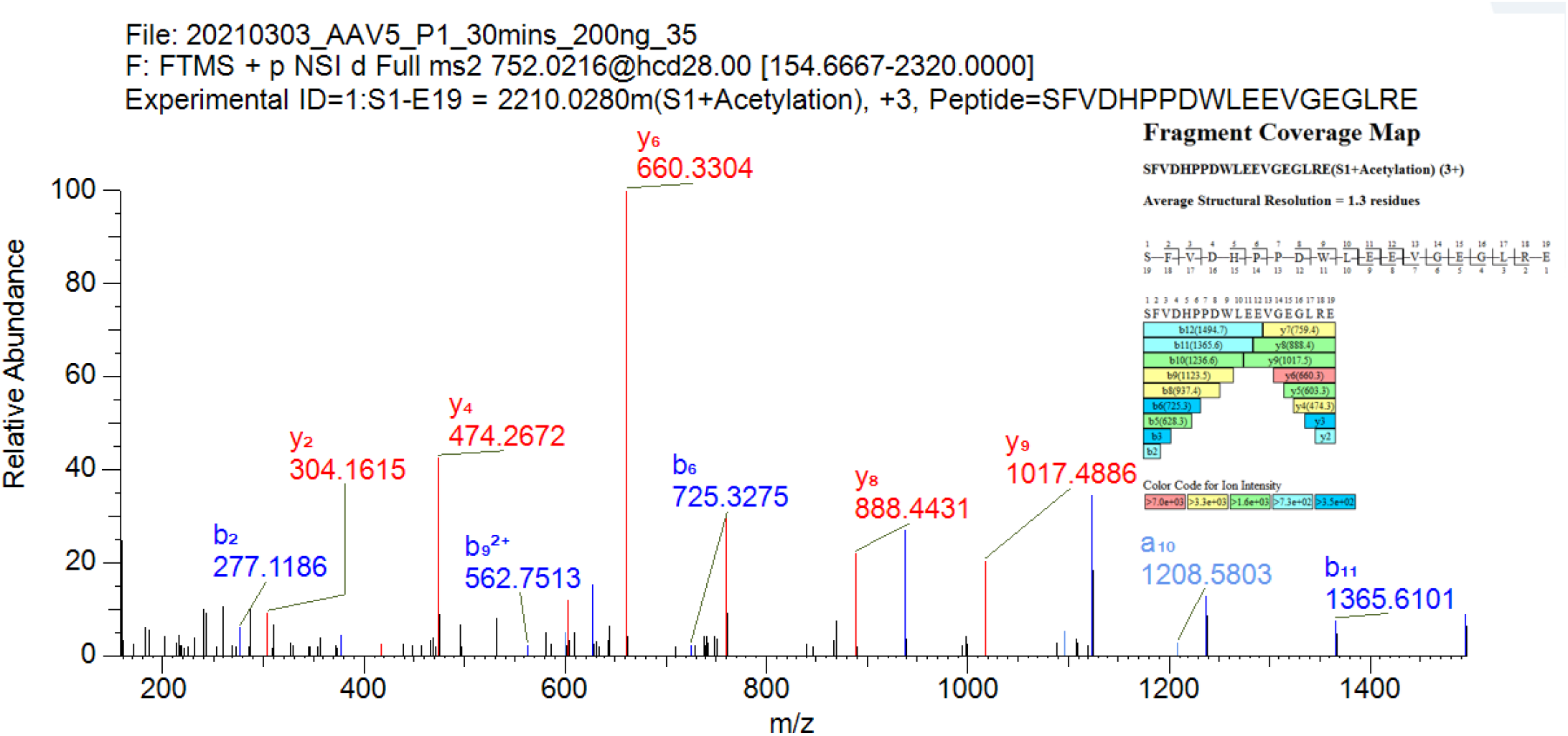
MS/MS spectrum with peak assignments and fragment coverage map for component (SFVDHPPDWLEEVGEGLRE) from Table S2 (S1+Acetylation from figure 3) using BioPharma Finder software.

**Figure S3:**
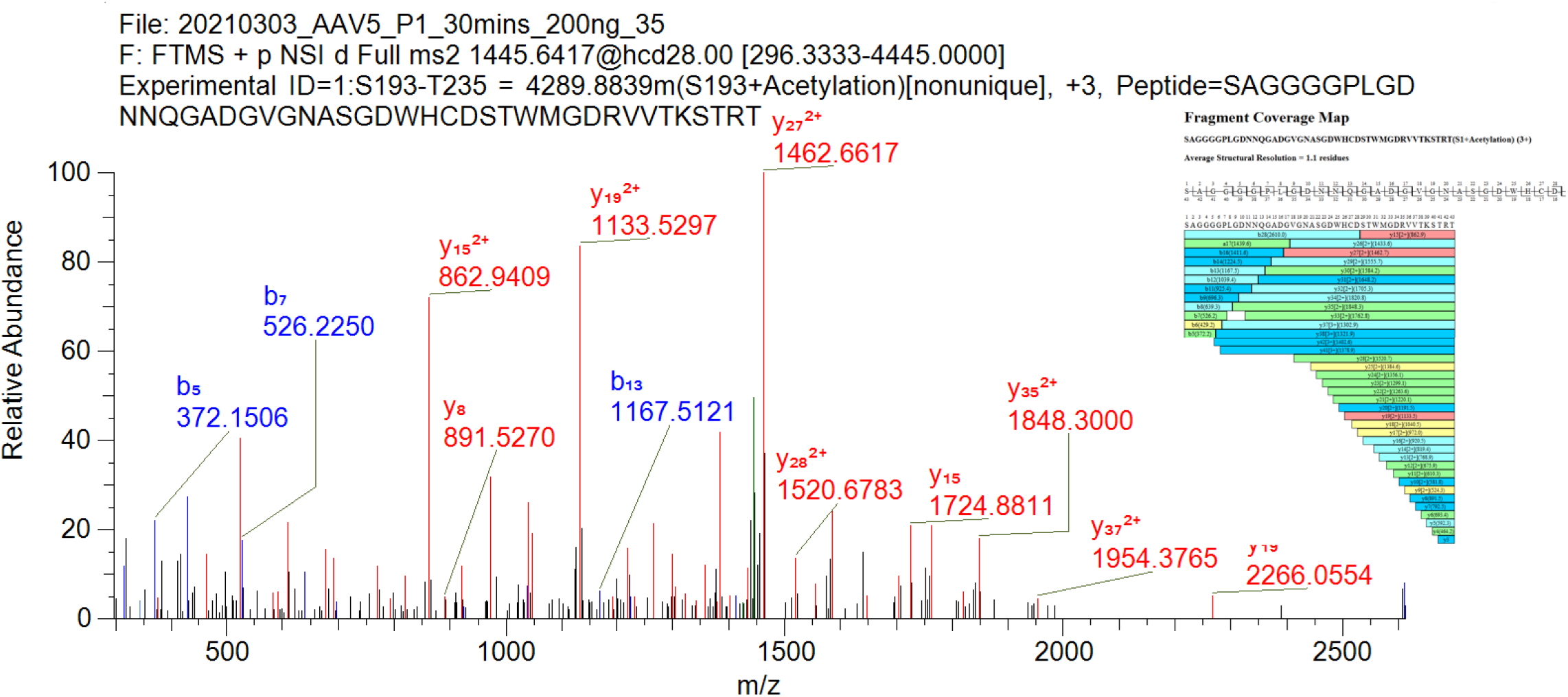
MS/MS spectrum with peak assignments and fragment coverage map for component (SAGGGGPLGDNNQGADGVGNASGDWHCDSTWMGDRVVTKSTRT) from Table S2 (S193+Acetylation from figure 3) using BioPharma Finder software.

**Figure S4:**
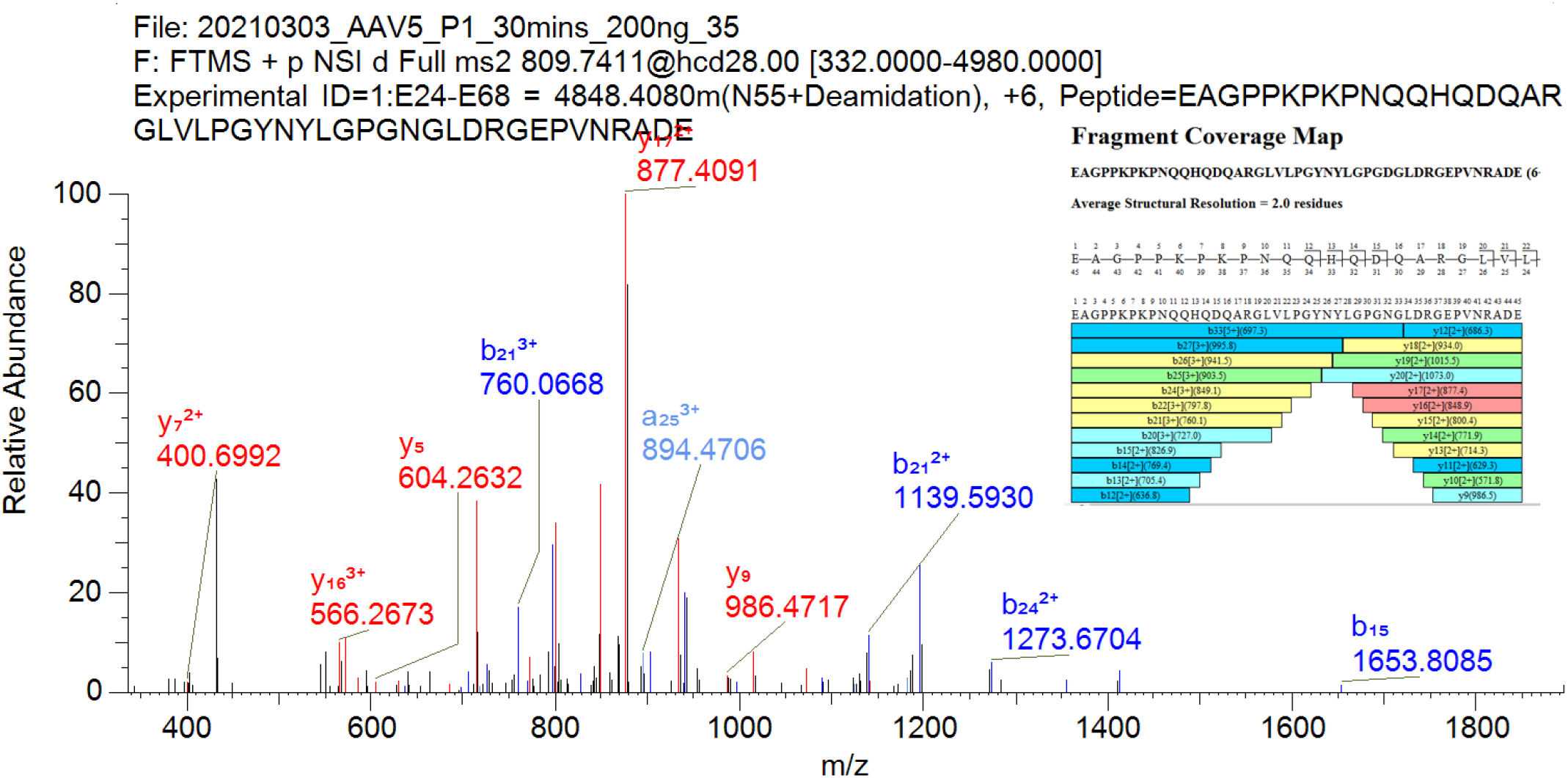
MS/MS spectrum with peak assignments and fragment coverage map for component (EAGPPKPKPNQQHQDQARGLVLPGYNYLGPGNGLDRGEPVNRADE) from Table S2 (N55+Deamidation from figure 3) using BioPharma Finder software.

**Figure S5:**
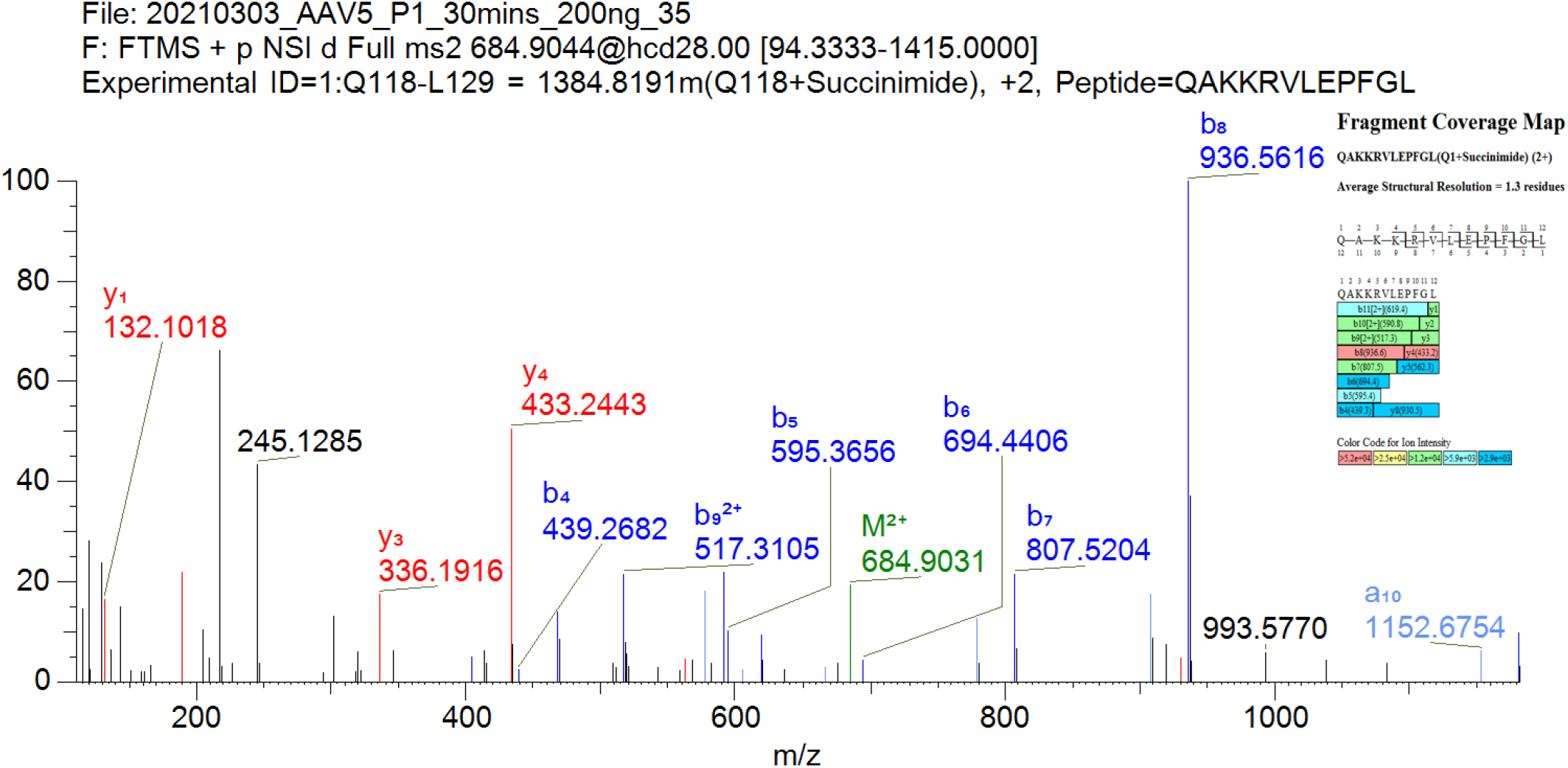
MS/MS spectrum with peak assignments and fragment coverage map for component (QAKKRVLEPFGL) from Table S2 (Q118+ Succinimide from figure 3) using BioPharma Finder software.

**Figure S6:**
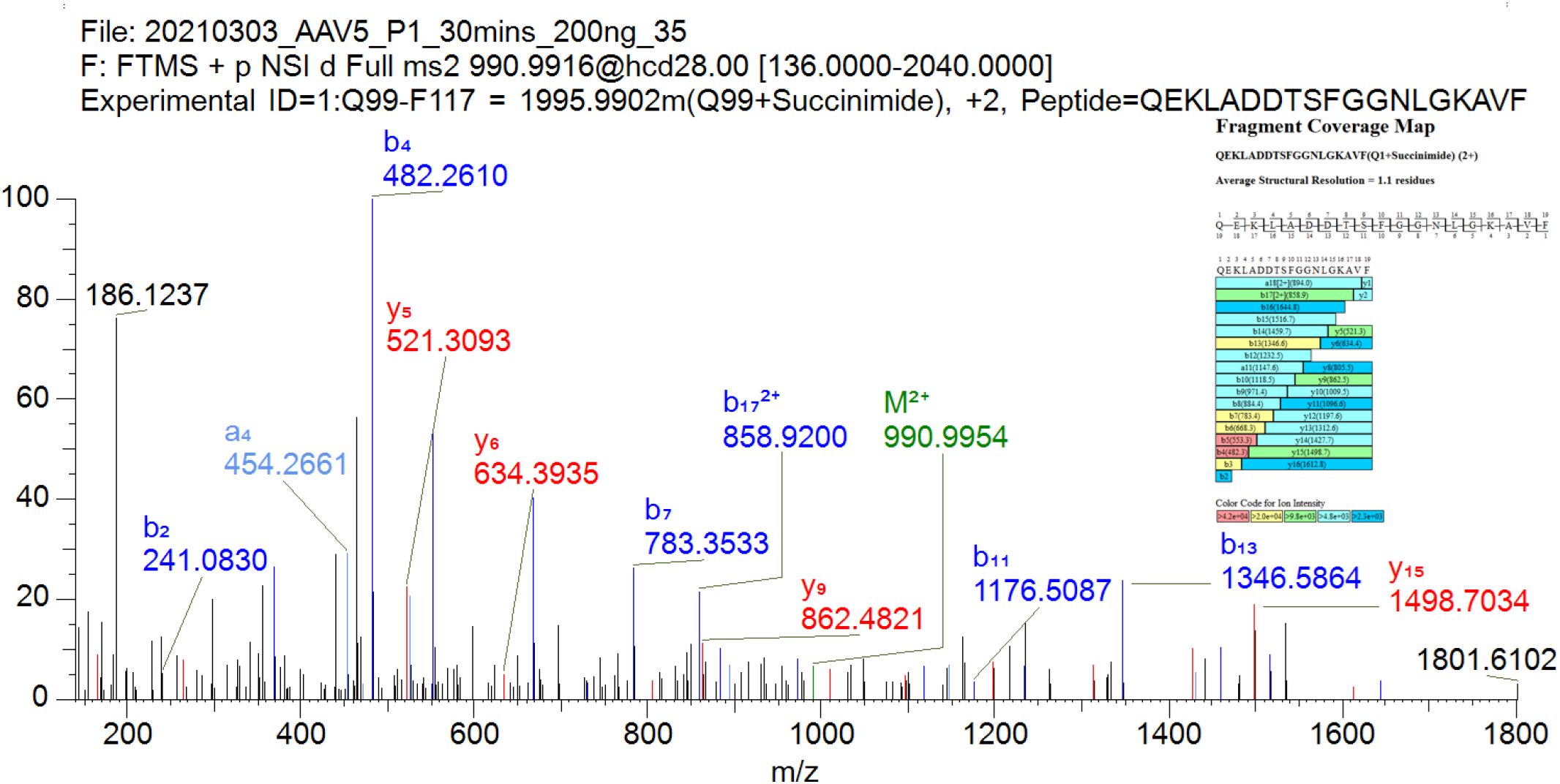
MS/MS spectrum with peak assignments and fragment coverage map for component (QEKLADDTSFGGNLGKAVF) from Table S2 (Q99+ Succinimide from figure 3) using BioPharma Finder software.

## References

[1] P.D. Kessler, G.M. Podsakoff, X. Chen, S.A. McQuiston, P.C. Colosi, L.A. Matelis, G.J. Kurtzman, B.J. Byrne, Gene delivery to skeletal muscle results in sustained expression and systemic delivery of a therapeutic protein, Proc. Natl. Acad. Sci. U. S. A. 93 (1996) 14082–14087. https://doi.org/10.1073/pnas.93.24.14082.

[2] X. Xiao, J. Li, R.J. Samulski, Efficient long-term gene transfer into muscle tissue of immunocompetent mice by adeno-associated virus vector., J. Virol. 70 (1996) 8098–8108. https://doi.org/10.1128/jvi.70.11.8098-8108.1996.

[3] D. Wang, P.W.L. Tai, G. Gao, Adeno-associated virus vector as a platform for gene therapy delivery, Nat. Rev. Drug Discov. 18 (2019) 358–378. https://doi.org/10.1038/s41573-019-0012-9.

[4] X. Jin, L. Liu, S. Nass, C. O’Riordan, E. Pastor, X.K. Zhang, Direct Liquid Chromatography/Mass Spectrometry Analysis for Complete Characterization of Recombinant Adeno-Associated Virus Capsid Proteins, Hum. Gene Ther. Methods. 28 (2017) 255–267. https://doi.org/10.1089/hgtb.2016.178.

[5] M. Nonnenmacher, T. Weber, Intracellular transport of recombinant adeno-associated virus vectors, Gene Ther. 19 (2012) 649–658. https://doi.org/10.1038/gt.2012.6.

[6] B. Mary, S. Maurya, S. Arumugam, V. Kumar, G.R. Jayandharan, Post-translational modifications in capsid proteins of recombinant adeno-associated virus (AAV) 1-rh10 serotypes, FEBS J. 286 (2019) 4964–4981. https://doi.org/10.1111/febs.15013.

[7] A. Bennett, S. Patel, M. Mietzsch, A. Jose, B. Lins-Austin, J.C. Yu, B. Bothner, R. McKenna, M. Agbandje-McKenna, Thermal Stability as a Determinant of AAV Serotype Identity, Mol. Ther. - Methods Clin. Dev. 6 (2017) 171–182. https://doi.org/10.1016/j.omtm.2017.07.003.

[8] B. Lins-Austin, S. Patel, M. Mietzsch, D. Brooke, A. Bennett, B. Venkatakrishnan, K. Van Vliet, A.N. Smith, J.R. Long, R. McKenna, M. Potter, B. Byrne, S.L. Boye, B. Bothner, R. Heilbronn, M. Agbandje-McKenna, Adeno-associated virus (AAV) capsid stability and liposome remodeling during endo/lysosomal pH trafficking, Viruses. 12 (2020). https://doi.org/10.3390/v12060668.

[9] Y. Xu, P. Guo, J. Zhang, M. Chrzanowski, H. Chew, J.A. Firrman, N. Sang, Y. Diao, W. Xiao, Effects of Thermally Induced Configuration Changes on rAAV Genome’s Enzymatic Accessibility, Mol. Ther. - Methods Clin. Dev. 18 (2020) 328–334. https://doi.org/10.1016/j.omtm.2020.06.005.

[10] L. Gigout, P. Rebollo, N. Clement, K.H. Warrington, N. Muzyczka, R.M. Linden, T. Weber, Altering AAV tropism with mosaic viral capsids, Mol. Ther. 11 (2005) 856–865. https://doi.org/10.1016/j.ymthe.2005.03.005.

[11] D. Rathore, A. Faustino, J. Schiel, E. Pang, M. Boyne, S. Rogstad, The role of mass spectrometry in the characterization of biologic protein products, Expert Rev. Proteomics. 15 (2018) 431–449. https://doi.org/10.1080/14789450.2018.1469982.

[12] S. Millán-Martín, C. Jakes, S. Carillo, T. Buchanan, M. Guender, D.B. Kristensen, T.M. Sloth, M. Ørgaard, K. Cook, J. Bones, Inter-laboratory study of an optimised peptide mapping workflow using automated trypsin digestion for monitoring monoclonal antibody product quality attributes, Anal. Bioanal. Chem. 412 (2020) 6833–6848. https://doi.org/10.1007/s00216-020-02809-z.

[13] J.C. Powers, A.D. Harley, D. V. Myers, Subsite specificity of porcine pepsin., Adv. Exp. Med. Biol. 95 (1977) 141–157. https://doi.org/10.1007/978-1-4757-0719-9_9.

[14] J. Ahn, M.J. Cao, Y.Q. Yu, J.R. Engen, Accessing the reproducibility and specificity of pepsin and other aspartic proteases, Biochim. Biophys. Acta - Proteins Proteomics. 1834 (2013) 1222–1229. https://doi.org/10.1016/j.bbapap.2012.10.003.

[15] N.G. Rumachik, S.A. Malaker, N.K. Paulk, VectorMOD: Method for Bottom-Up Proteomic Characterization of rAAV Capsid Post-Translational Modifications and Vector Impurities, Front. Immunol. 12 (2021) 830. https://doi.org/10.3389/fimmu.2021.657795.

[16] D.X. Zhang, D.X. Jin, D.L. Liu, D.Z. Zhang, D.S. Koza, D.Y.Q. Yu, D.W. Chen, Optimized reversed phase LC/MS methods for intact protein analysis and peptide mapping of adeno-associated virus (AAV) proteins, Https://Home.Liebertpub.Com/Hum. (2021). https://doi.org/10.1089/HUM.2021.046.

[17] A. Bennett, S. Patel, M. Mietzsch, A. Jose, B. Lins-Austin, J.C. Yu, B. Bothner, R. McKenna, M. Agbandje-McKenna, Thermal Stability as a Determinant of AAV Serotype Identity, Mol. Ther. - Methods Clin. Dev. 6 (2017) 171–182. https://doi.org/10.1016/j.omtm.2017.07.003.

[18] S. Liu, K.R. Moulton, J.R. Auclair, Z.S. Zhou, Mildly acidic conditions eliminate deamidation artifact during proteolysis: Digestion with endoprotease Glu-C at pH 4.5, Amino Acids. 48 (2016) 1059–1067. https://doi.org/10.1007/s00726-015-2166-z.

[19] M. Haberger, A.K. Heidenreich, T. Schlothauer, M. Hook, J. Gassner, K. Bomans, M. Yegres, A. Zwick, B. Zimmermann, H. Wegele, L. Bonnington, D. Reusch, P. Bulau, Functional assessment of antibody oxidation by native mass spectrometry, MAbs. 7 (2015) 891–900. https://doi.org/10.1080/19420862.2015.1052199.

[20] J. Cacia, R. Keck, L.G. Presta, J. Frenz, Isomerization of an aspartic acid residue in the complementarity-determining regions of a recombinant antibody to human IgE: Identification and effect on binding affinity, Biochemistry. 35 (1996) 1897–1903. https://doi.org/10.1021/bi951526c.

[21] M. Cao, W. Xu, B. Niu, I. Kabundi, H. Luo, M. Prophet, W. Chen, D. Liu, S. V. Saveliev, M. Urh, J. Wang, An Automated and Qualified Platform Method for Site-Specific Succinimide and Deamidation Quantitation Using Low-pH Peptide Mapping, J. Pharm. Sci. 108 (2019) 3540–3549. https://doi.org/10.1016/j.xphs.2019.07.019.

